# Impact of Soil Salinity on Microbial Community Composition in Coastal Agricultural lands of Bangladesh: A Metagenomic Approach

**DOI:** 10.1101/2025.02.08.637242

**Authors:** Sumit Das, Maksudur Rahman Nayem, Kamrun Nahar Hira, Romana Shermin, Md. Razib Hosen, Md. Amzad Hossain, Md. Tariquzzaman, Razib biswas, Farazi Abinash Rahman, Mithun Kumar Saha, Mohammad Fazle Alam Rabbi

## Abstract

A significant obstacle to agricultural productivity is soil salinity, especially in coastal areas where the problem is made worse by climate change and rising sea levels. This study investigates the impact of salinity on microbial community structure and function in agricultural soils from Kuakata, Bangladesh, using 16S rRNA metagenomic sequencing technique. Both saline (EC > 4 dS/m) and non-saline (EC < 1.5 dS/m) areas had soil samples taken; saline soils showed symptoms of crop stress, such as yellowing and blackish discoloration. In comparison to their non-saline counterparts, our investigation showed that saline soils had much reduced microbial diversity and different community compositions. Both soil types were dominated by Proteobacteria and Chloroflexi, while saline soils displayed an enrichment of anaerobic texa (e.g., Desulfobacterota, Pseudomonas) and sulfate-reducing bacteria (e.g., Desulfobacterota, Deferrisomatota). Interestingly, saline soils had a much larger percentage of anaerobic microorganisms (36.4%) than non-saline soils (29.3%) which might results in enhanced denitrification, sulphate reduction, and hydrogen sulphide buildup. These processes disrupt nutrient cycling, lower organic matter decomposition and create toxic conditions for plant growth, ultimately contributing to crop failure. This study provides an insight into the distinct microbial composition in saline and non-saline samples, which can assist in understanding the role of microbial populations in agriculture in saline environments.

## 1. Introduction

Soil salinity is a significant global issue, affecting approximately 20% of the world’s cultivated lands ^1^. This problem is expected to worsen in the coming decades, with projections indicating that up to 50% of arable lands could be impacted by salinity by 2050 ^2^. Such a substantial loss of farmland would severely reduce crop yields, directly contradicting the growing global demand for food and fiber.

Soil salinity is characterized by the accumulation of excessive ions, such as calcium (Ca²⁺), magnesium (Mg²⁺), sodium (Na⁺), sulfates (SO₄²⁻), and chlorides (Cl⁻), which hinder plant growth and function ^3^. Climate change, particularly sea level rise due to global warming, is increasingly contributing to soil salinity ^4^.

Microorganisms are essential components of the soil ecosystem, influencing its multidimensional functionality ^5^. They play critical roles in processes such as oxidation, nitrification, ammonification, nitrogen fixation, and the decomposition of organic matter, which facilitate nutrient transformation ^6^. Additionally, microbes store carbon and nutrients in their biomass, releasing them into the soil ^7^. They are also vital for plant health, as they associate with roots and solubilize essential nutrients for plant uptake ^8^. However, microbial communities are sensitive to changes in soil conditions, including salinity, which significantly impacts their composition, diversity, and function ^9^. High salt concentrations affect microbes through osmotic stress and specific ion effects ^10^.

Advances in molecular techniques, particularly next-generation sequencing, have made metagenomics a powerful tool for studying microbial communities. Metagenomic sequencing provides an efficient way to analyze changes in microbial diversity and community structure under salt stress ^11^.

In Bangladesh, soil salinity is a growing concern, particularly in coastal regions. The country has a total land area of 147,570 square kilometers, with 29,000 square kilometers located in coastal areas ^12^. According to the Soil Resources Development Institute (SRDI, 2010), the area affected by salinity increased from 83.3 million hectares in 1973 to 102 million hectares in 2000, and further to 105.6 million hectares by 2009. This represents a significant portion of the country’s arable land, posing a threat to crop production and the economy.

Given the critical role of microbes in soil health and plant growth, understanding their behavior in saline environments is essential. In this study, we employed 16S metagenomic sequencing to investigate the rhizosphere microbiomes of crop fields in both saline and non-saline soils from the coastal region of Bangladesh. This research aims to provide insights into the microbial diversity differences between saline and non-saline soils and identify specific microbes that are essential for healthy crop growth.

## **2.** Materials and methods

### 2.1 Soil sample collection and preparation

The soil samples were collected from 6 locations around the coastal region of Kuakata. Among these locations, 3 were selected for the collection of 10 saline soil samples and 3 were selected for the collection of 6 non-saline soil samples. From the non-saline group, two samples (NS1, NS2) were replicated for the metagenomics study along with other samples (n=18). The saline soils are taken from agricultural land in the coastal zone where the entire crop yield was lost due to salinity. The crops turned yellow to blackish in appearance and a visible layer of salt precipitate was present in the field. The names, GPS coordinates (collected using Google Maps), and corresponding soil samples of the locations are given in the supplementary Table 1. The sampling points are also given in the Figure 1. The main parameter used to differentiate saline soils from non-saline soils was the Electrical Conductivity (EC) of the soils. The EC of the soils was measured using a hand-held Soil EC meter. Soils with EC greater than 4 dSm^-1^ were selected as saline soil samples and soils with EC less than 1.5 dSm^-1^ were selected as non-saline soil samples. The rhizobium soil samples were gathered randomly using a composite sampling technique from a depth of approximately 10 to 15 cm or 6 inches, near agricultural fields. Each soil samples were collected in 50 ml sterile falcon tubes for metagenomic analysis and in plastic bags (1 kg) for soil physicochemical and chemical analysis. The soil samples collected in the falcon tubes were transported to the Next Generation Sequencing Laboratory maintaining the cold chain and stored at -80°C until DNA extraction was executed. The soil samples collected for soil physicochemical and chemical analyses were subsequently air-dried and rid of any visible roots or debris. To expedite the drying process sunlight exposure was employed. Following this, the larger soil fragments were delicately fragmented using a wooden hammer and sifted through a stainless-steel sieve of 2 mm mesh size. These sifted samples were then placed into plastic containers, appropriately labeled, and set aside for subsequent physicochemical and chemical analyses.

**Figure 1:**
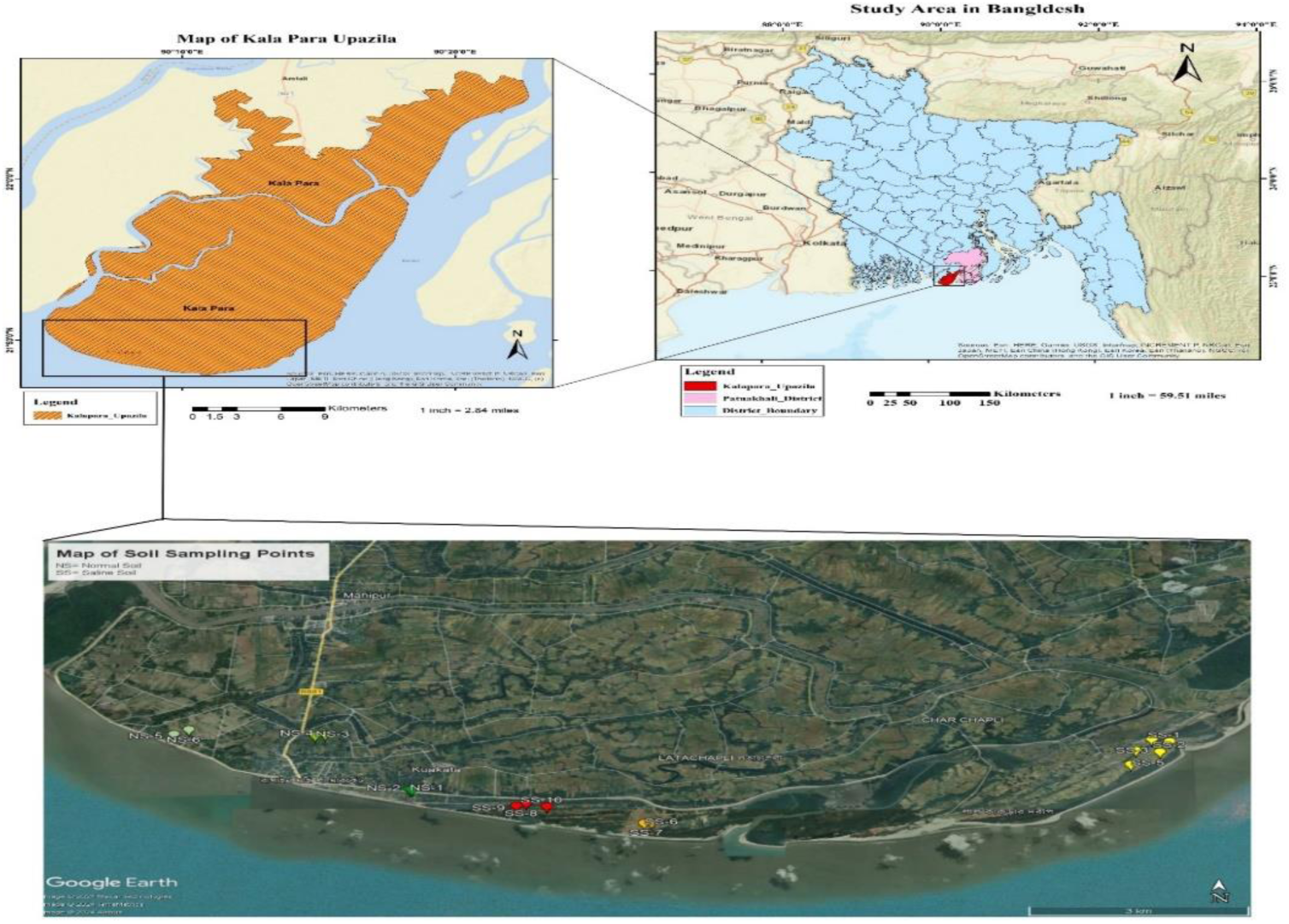
Map of the study area and soil sampling points.

### 2.2 DNA extraction from soil samples for molecular processing

The DNA extraction of the collected soil samples was carried out using the Invitrogen™ PureLink™ Microbiome DNA Purification kit by ThermoFisher Scientific, USA. The purified DNA extracted samples were quantified using a Qubit 4 and NanoDrop2000 (Thermo Scientific, USA) to determine the concentration and relative purities by A260/280 ratio, before starting the library preparation for 16S metagenomics.

### 2.3 Library preparation and sequencing

The V1-V9 hypervariable region of the 16S rRNA gene was amplified using the 27F and 1492R primer pair. Polymerase chain reaction (PCR) was performed using Q5® High-Fidelity 2X Master Mix following the manufacturer’s guidelines. The thermal cycling protocol consisted of an initial denaturation step at 95°C for 3 minutes, followed by 30 cycles of denaturation at 95°C for 15 seconds, annealing at 60°C for 15 seconds, and extension at 72°C for 80 seconds, with a final elongation step at 72°C for 10 minutes. Post-amplification, the DNA was purified using AMPure XP reagent (Beckman Coulter, Inc.) and quantified using the Qubit 4 fluorometer with Qubit™ 1X dsDNA High Sensitivity (HS) reagent (Thermo Fisher Scientific, USA).

For library preparation, 200 ng of purified DNA was processed for end repair and dA-tailing, followed by barcode ligation using the Native Barcoding Kit 24 V14 (SQK-NBD114.24) from Oxford Nanopore. The barcoded libraries were pooled, and sequencing adapters were ligated using the NEBNext® Quick Ligation Module (NEB, E6056). The adapter-ligated library were loaded onto the R10.4.1 flow cell (FLO-MIN114; Oxford Nanopore Technologies). Sequencing was conducted on the MinION™ Mk1C device, with data acquisition performed using MINKNOW software version 1.11.5 (Oxford Nanopore Technologies).

### 2.4 Bioinformatics analysis

Basecalling of the sequencing data (FAST5 files) was performed using Guppy version 6.3.2 (Oxford Nanopore Technologies) in high-accuracy mode, generating pass reads in FASTQ format with a minimum quality score greater than 9. Adapter and barcode sequences were removed using Porechop v0.2.4, and low-quality bases were trimmed using Nanofilt. The quality-trimmed reads were filtered by size, retaining sequences above 1400 base pairs (bp) to match the expected size distribution of the V1-V9 region of the 16S rRNA gene. Taxonomic classification of the reads was conducted using the wf-16s workflow in Epi2me software (Oxford Nanopore Technologies), with SILVA as the reference database and Kraken2 as the classifier. All other parameters were maintained at their default settings.

### 2.5 Statistical analysis

Downstream analysis, including alpha and beta diversity, microbial composition, differential abundance calculation, and statistical comparisons, was performed using the Phyloseq (v4.2), Vegan, ggplot2, ggpubr, and microbiomeutilities packages in R (v4.3.0). Data filtration was then carried out in a manner that at least 5 OTUs was required to be present for any specific genre for any specific sample. Filtered data was normalized using the Total Sum Scaling method, and the normalized data were utilized to calculate alpha and beta diversity. Alpha diversity, representing within-sample diversity, was assessed using the observed OTUs, Chao1, ACE, Shannon, Simpson, and Inverse Simpson indices, computed with the Phyloseq and Vegan packages. Differences in microbial diversity and abundance between groups were evaluated using the Wilcoxon rank-sum test from the microbiomeutilities R package.

To examine differences in microbial diversity between sample groups (β-diversity), Principal Coordinate Analysis (PCoA) based on the Jaccard distance matric was performed. Permutational Multivariate Analysis of Variance (PERMANOVA) with 999 permutations was used to determine statistical significance (p-value) for group differences. Functional annotation prediction was conducted using the Metagenassist tool.

## 3. Results

### 3.1 Physicochemical properties of soil samples

A total 16 samples were gathered from coastal areas. 8 of them were non-saline, and 10 were saline samples. With a p-value of 0.00044, saline soils exhibit a substantially higher electrical conductivity (median of 4.82 dS/m) than non-saline soils (median of 0.5 dS/m).

In the case of ESP, the proportion of exchangeable sodium in saline soils was higher (median 33.78%) than in non-saline soils (median 11.37%), indicating a significant difference (p = 0.0016). With a p-value of 0.011, saline soils had a substantially higher pH (median 6.83) than non-saline soils (median 5.40). The cation exchange capacity of saline soils was significantly larger (median about 16.08 meq/100g) than that of non-saline soils (median 3.35 meq/100g), with p = 0.00044. A significant difference was observed (p = 0.00044) with saline soils exhibiting a much higher median sulfur content (288.9mg/kg) compared to non-saline soils (120.62). There was a significant difference (p = 0.00044) with saline soils having a higher sodium adsorption ratio (median 2.38) compared to non-saline soils (median 0.41).

The comparison of the physicochemical and chemical parameters between non-saline soils and saline soils is shown in Figure 2.

**Figure 2:**
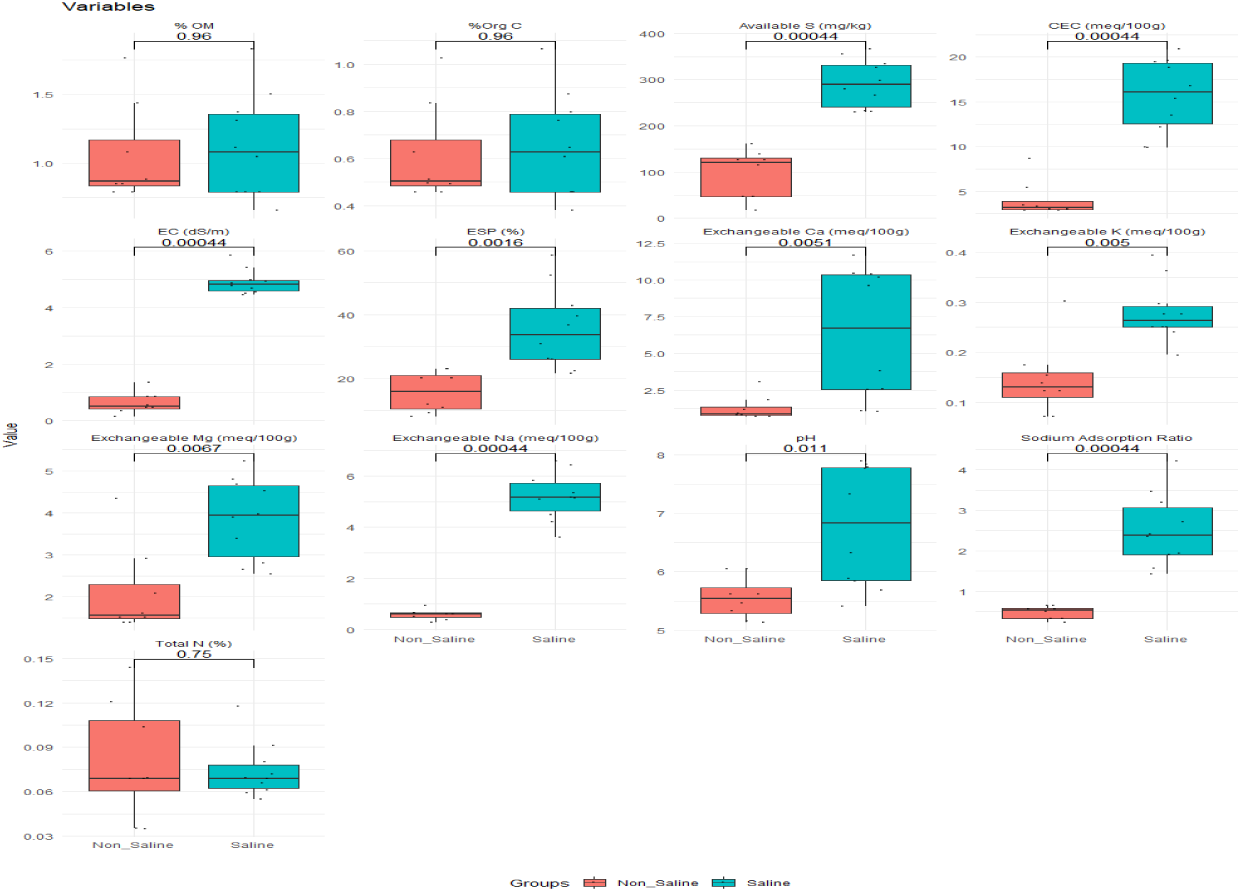
Boxplots were used to plot the physicochemical parameters, and the Wilcoxon rank sum test was used for comparisons. The significance level (*p*-value) was calculated using the Wilcoxon rank sum test.

### 3.2 Metagenomic analyses and microbiota profiles of soil samples

#### 3.2.1: Multivariate analysis and Taxonomic clustering

From the sequencing data, Epi2me generated a total of 1901 operational taxonomic units. After filtering out, 520 OTUs were remaining in the data set. From those OTUs, 153 were unique to saline samples and 106 were unique to non-saline samples. 261 OTUs were found to be shared by both of the groups. The distribution of OTUs are presented in figure 3.

**Figure 3:**
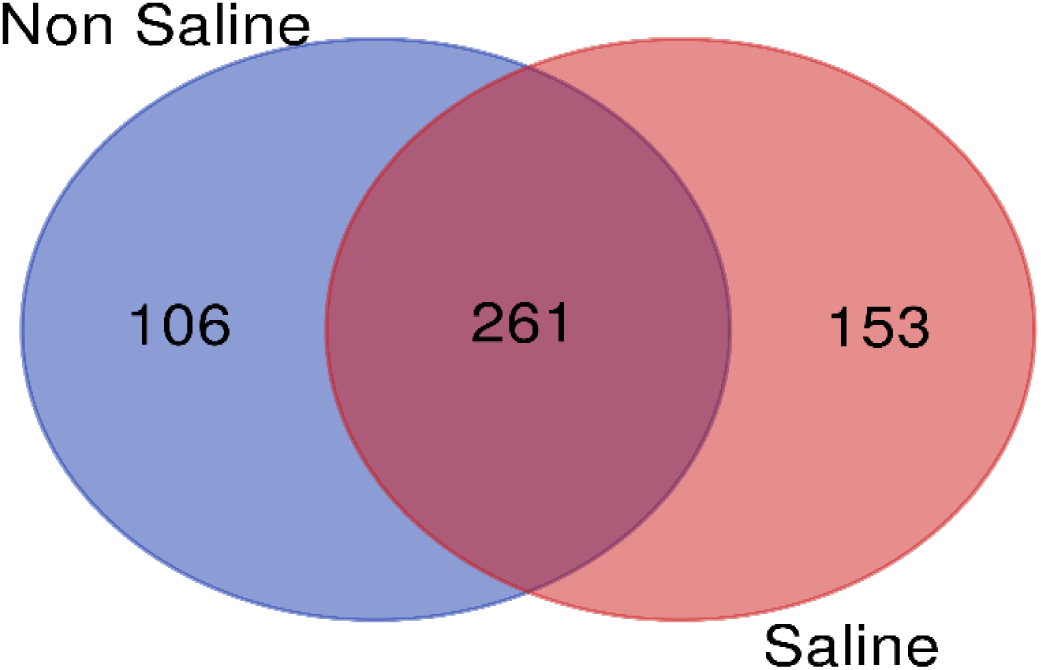
Van-diagram of OTU distribution between sample groups. Both unique and shared bacterial genera identified in the metagenomics study.

**Figure 4:**
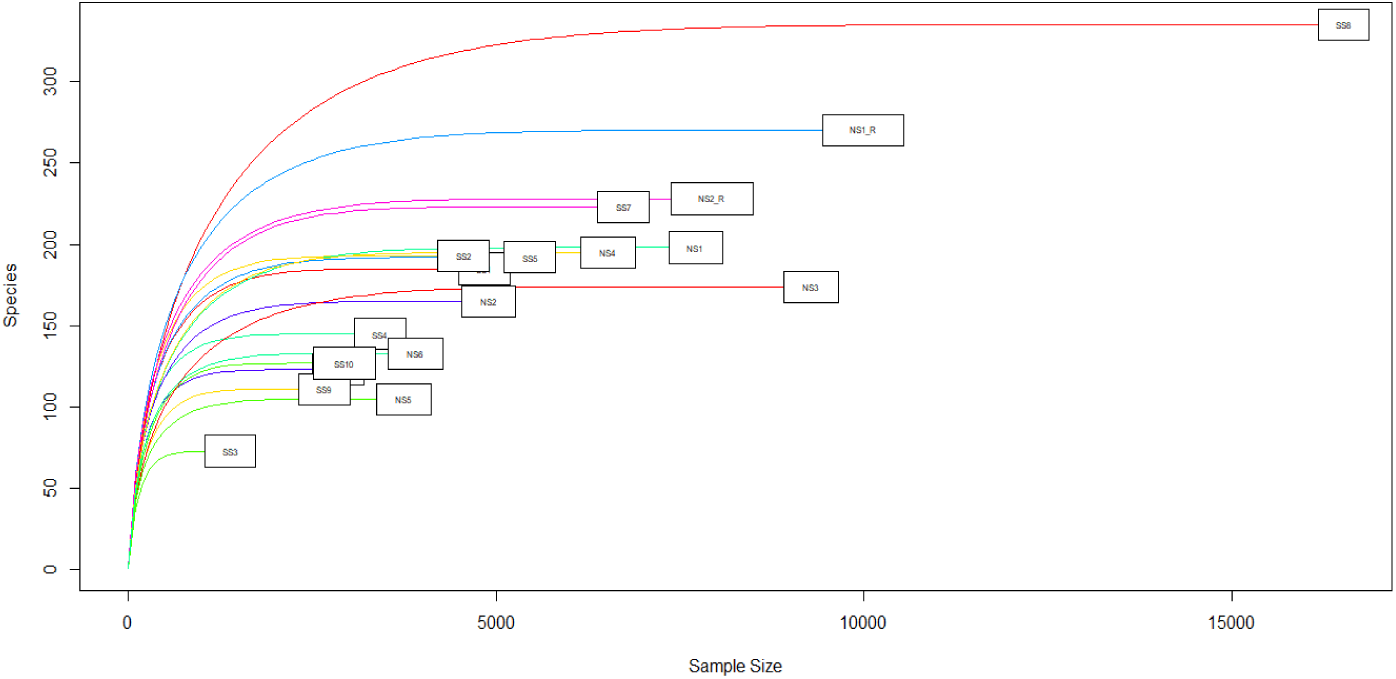
Rarefaction curve of the soil samples

A rarefaction curve was plotted to determine the number of OTUs against the number of reads which was needed for each soil sample. The rarefaction curve is shown in Figure 11. From the rarefaction curve, it can be seen that enough reads were generated for all of the samples to get the maximum number of OTUs from each specimen.

According to our analysis, the sequence data contained one archaeal phylum and a total of 26, bacterial phylum. There were 24 bacterial phyla in the group of saline samples and 22 bacterial phyla in the group of non-saline samples. Twenty of the phyla identified in this study were present in both sample groups. Four phyla (Calditrichota, Entotheonellaeota, Hydrogenedentes, and Modulibacteria) were exclusive to the saline group and not found in the non-saline sample, whereas two phyla (Armatimonadota and Campylobacterota) were only found in the non-saline sample. Although the Archeal phylum (Halobacterota) was present in both groups, non-saline samples had a significantly greater abundance (Fig-6A).

With a relative abundance of 29.9% for the non-saline sample group and 34.05% for the saline group, Proteobacteria was the most prevalent phylum in both the saline and non-saline sample groups. Chloroflexi came in second with relative abundances of 21.79% and 26.07% for the non-saline and saline groups, respectively. Desulfobacterota and Deferrisomatota, two common bacterial phyla in both groups, were significantly (P<0.05, Wilcoxon rank sum test) more prevalent in saline samples than in the non-saline group (Figure 6 A). However, the no-saline group had a noticeably higher (P<0.05, Wilcoxon rank sum test) abundance of Myxococcota, Methylomirabilota, Gemmatimonadota, Verrucomicrobiota, Spirochaetota, and Planctomycetota (Figure 6 A).

Figure 5 shows the taxonomic classification at the genus-level of microbiomes in saline and non-saline samples. The study found that 366 bacterial genera were present in all of the non-saline samples (Fig. 3). The top 10 most abundant genera were Anaerolineaceae_uncultured (13.3%), ADurb.Bin063-1 (4.8%), Anaerolinea (4.3%), Sideroxydans (2.7%), Candidatus Solibacter (2.6%), Candidatus Udaeobacter (2.5%), RBG-16-58-14 (2.1%), Geobacteraceae_uncultured (2%), Desulfobacca (1.9%), and Candidatus Nitrotoga (1.7%). However, saline samples contained a total of 414 bacterial taxa. Escherichia-Shigella (2%), RBG-16-58-14 (1.9%), Bacillus (1.7%), Thioalkalispira-Sulfurivermis (1.7%), Caldilineaceae_uncultured (1.5%), Anaerolineaceae_uncultured (21.4%), Pseudomonas (5.1%), Desulfobulbaceae_uncultured (3.4%), Desulfobacca (2.3%), and Escherichia-Shigella (2%) were top 10 genera in saline samples. The identified bacterial genera are clearly divided into two main clusters based on the kind of sample type in the heatmap depiction (Fig. 7) of the soil microbiomes.

**Figure 5:**
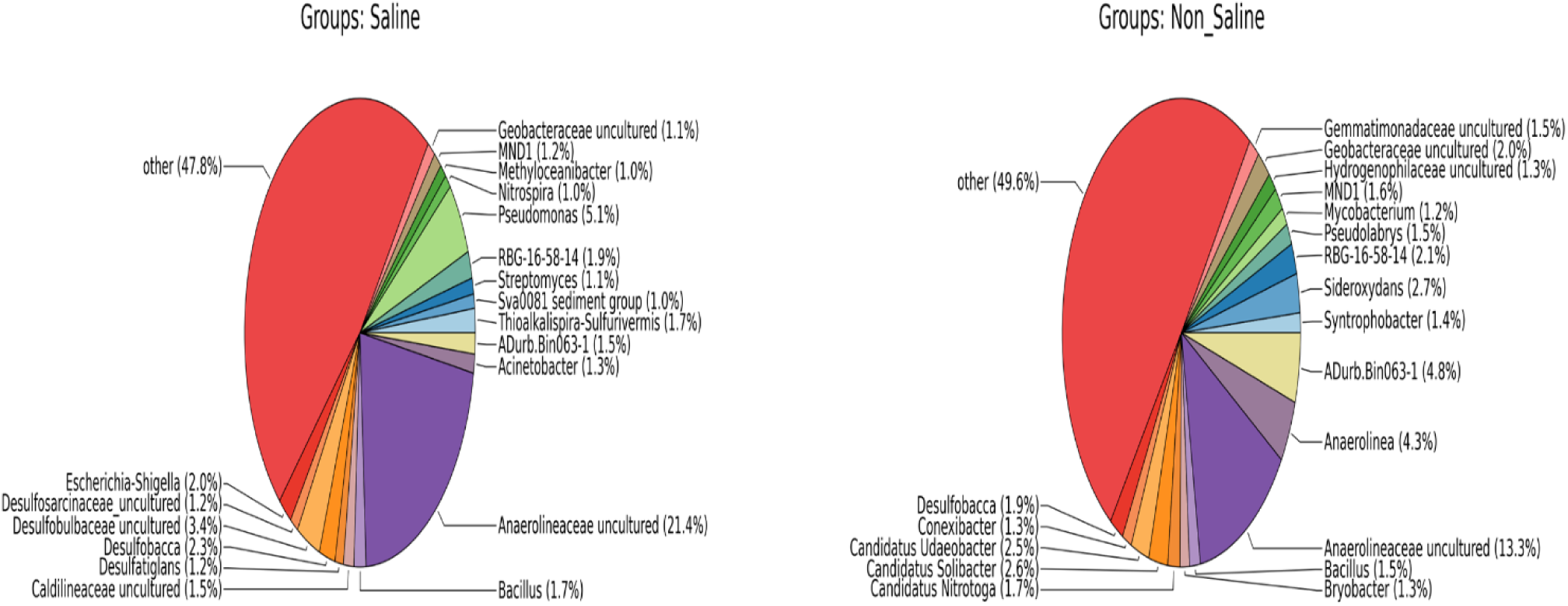
Taxonomic (Genus) abundance of different groups of microbiota in saline and non-saline samples

**Figure 6:**
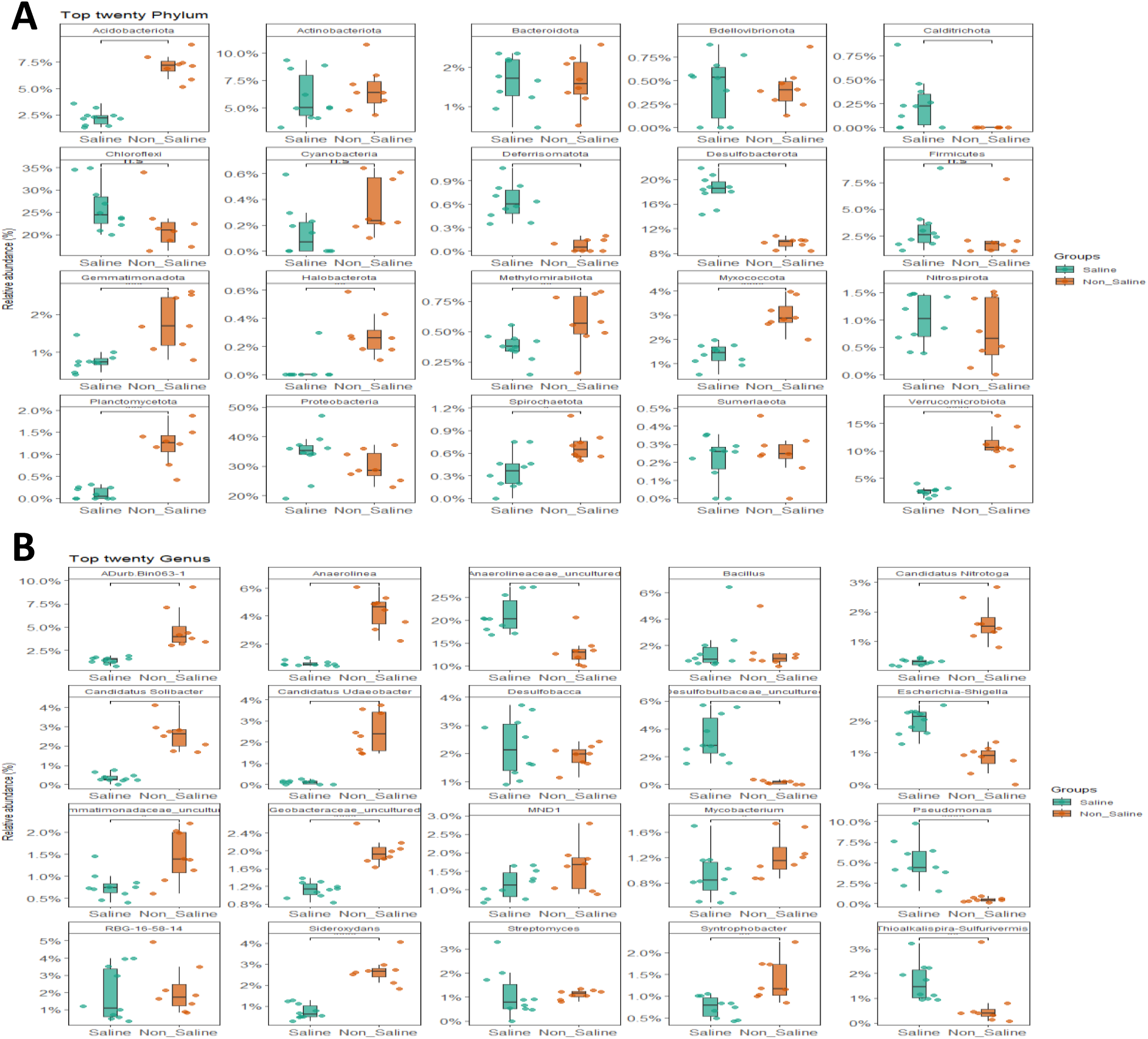
Top twenty bacterial phyla (A) and genus (B) in saline and non-saline samples. Significance level (*p* value, Wilcoxon rank sum test) 0.0001, 0.001, 0.01, 0.05, and 0.1 are represented by the symbols "****", "***", "**", "*", and "n.s" respectively.

**Figure 7:**
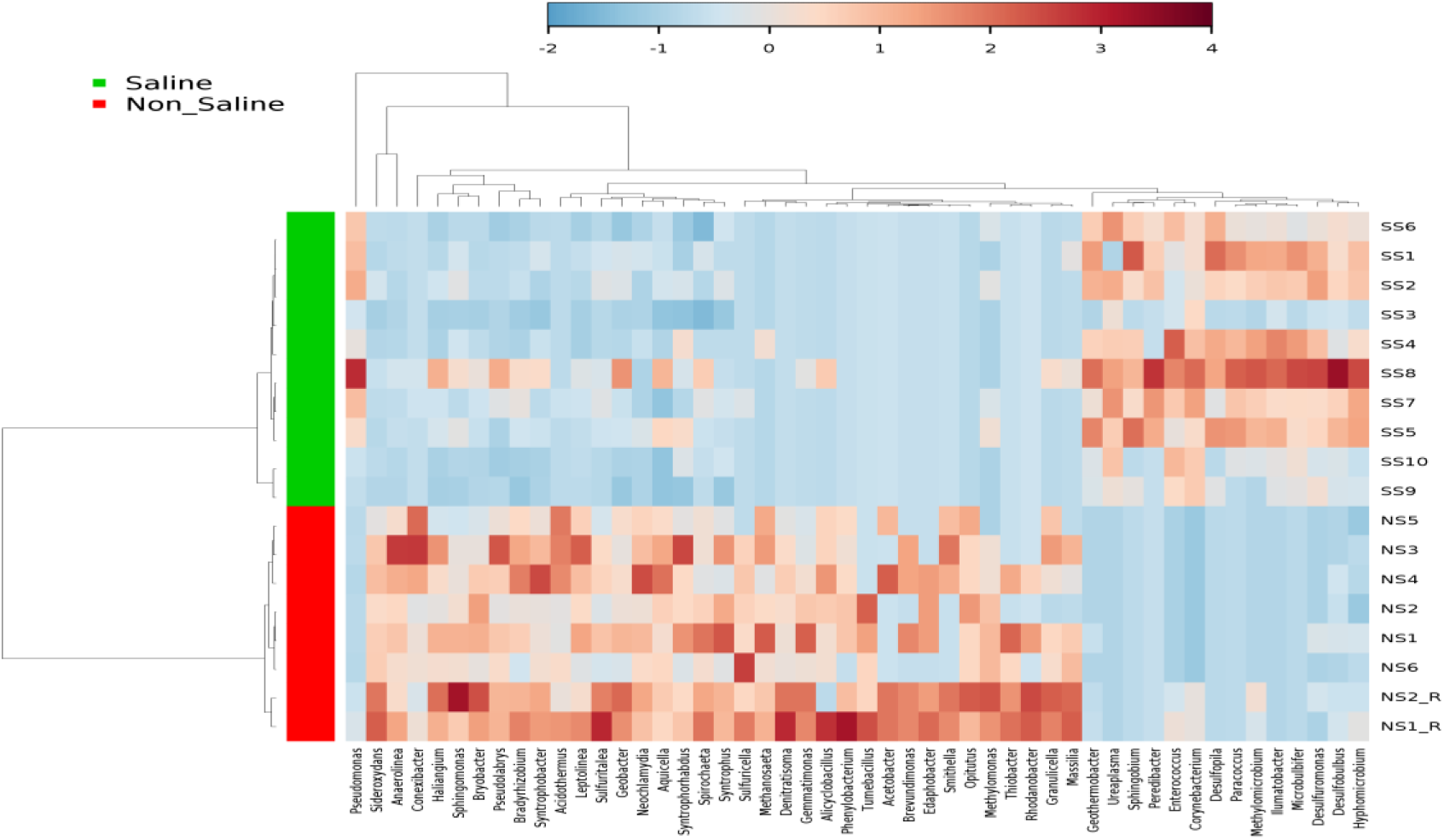
40 unique bacterial genera are displayed in a heatmap across sample categories according to the T-test. With Ward’s Minimum Variance Clustering Method, METAGENassist produced the heatmap. With a value ranging from -2 (lowest abundance) to 4 (highest abundance), the color codes show the prevalence and completeness of each genus in the relevant sample group. Blue cells represent less abundance, while red cells represent more abundant patterns.

#### 3.2.2 : Alpha-beta diversity and functional features analysis

The alpha-beta diversity of the microbiome in each kind of soil was examined to determine the differences in the microbiota of the groups. The Normalized data were examined for Observed, Chao1, Shannon, Simpson, and InvSimpson for Alpha diversity measurement. The result showed significant differences (P < 0.05, Kruskal-Wallis test and Wilcoxon signed rank test) in Simpson and InvSimpson indices in bacterial composition across the samples of both saline and non-saline groups. The observed species, Chao1, Shannon indices did not show statistically significant differences between the two conditions, though variability was higher in non-saline samples. Figure 8 shows the alpha diversity parameters of bacterial communities in saline and non-saline soil.

**Figure 8:**
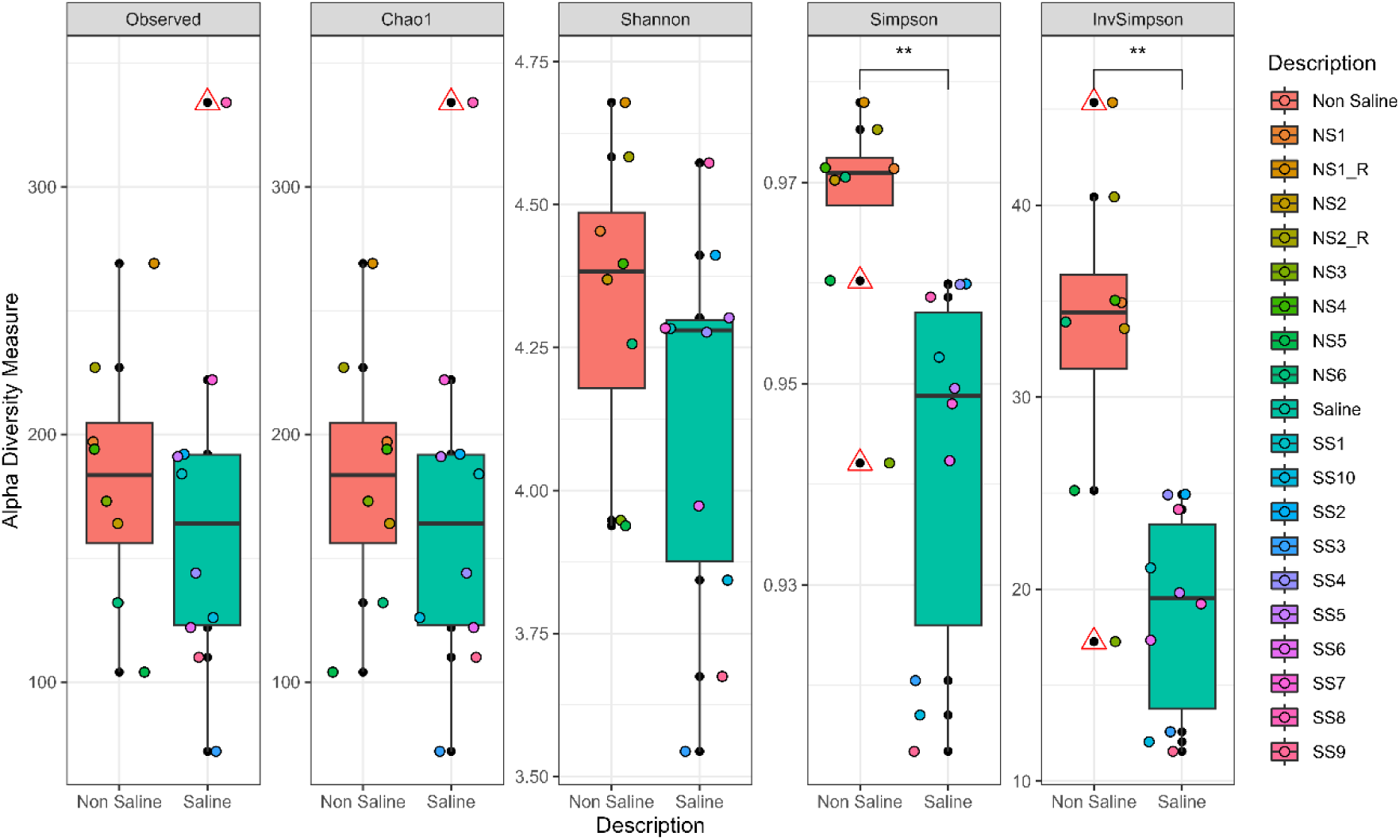
alpha diversity parameters of bacterial communities in saline and non-saline soil. Significance level (*p* value) 0.0001, 0.001, 0.01, 0.05, and 0.1 are represented by the symbols "****", "***", "**", "*", and "n.s" respectively.

There were significant variations between the two sample groups (beta diversity) according to principal coordinate analysis (PCoA) using Bray Curtis distance (Fig. 9A), and Jaccard distance (Fig. 9B) (PERMANOVA, p > 0.01). The non-metric multidimensional scaling (NMDS) method based on Bray Curtis distance and Jaccard distance produced similar results with significant variations (PERMANOVA, p > 0.01) (Fig. 9C, 9D). The stress value represents the goodness of fit of NMDS (> 0.2 Poor, 0.1–0.2, Fair, 0.05–0.1 Good, and < 0.05 Excellent) which was found to be 0.058.

**Figure 9:**
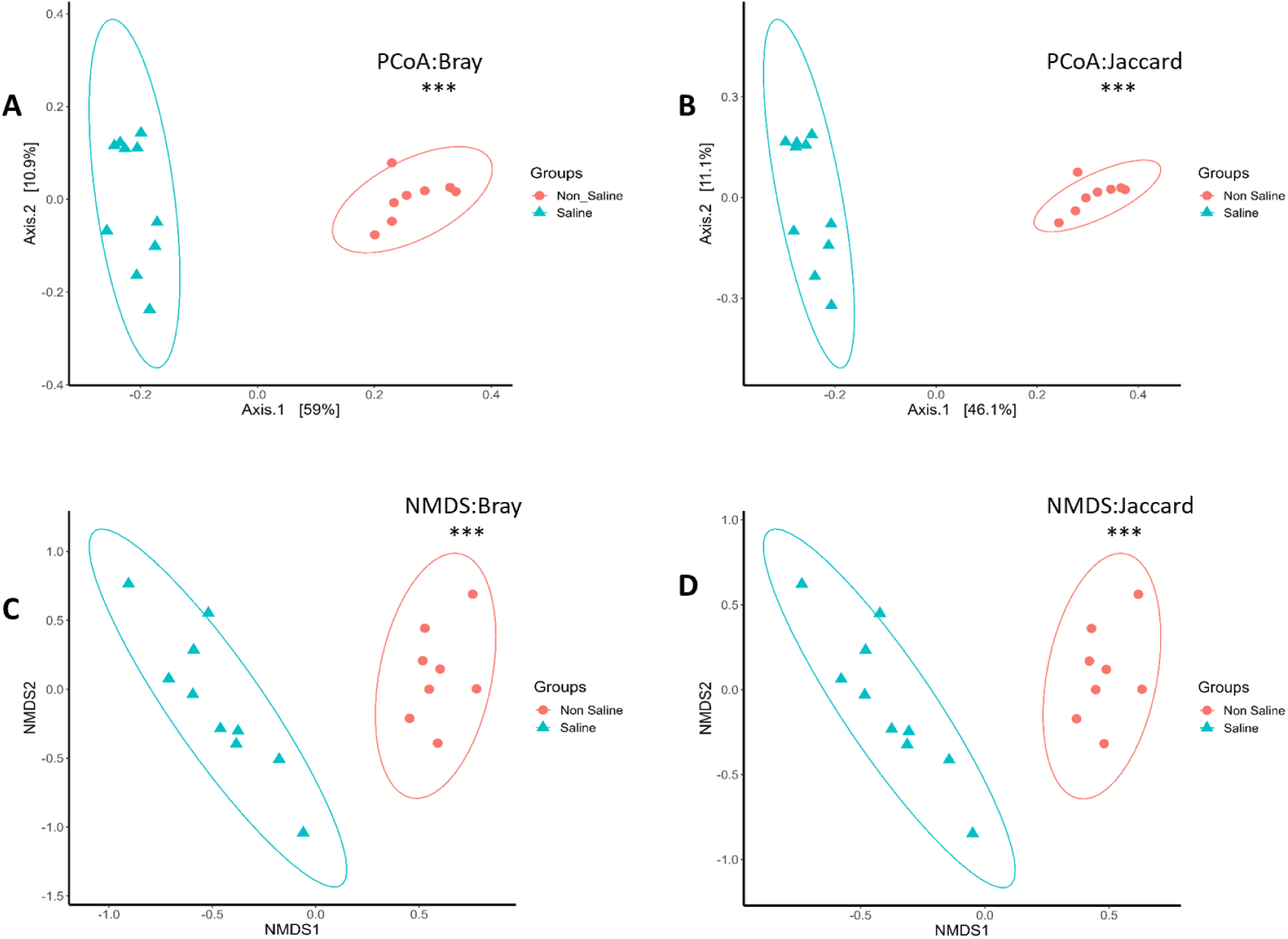
Beta diversity measures of microbial communities. Principal coordinate analysis (PCoA) (**A,B**) and non-metric multidimensional scaling (**C,D**) were performed using Bray and Jaccard distance metrics for the two groups of samples. PERMANOVA with 999 permutations was used to determine the significance (*p* value) of differences between two groups. Significance level (*p* value) 0.0001, 0.001, 0.01, 0.05, and 0.1 are represented by the symbols "****", "***", "**", "*", and “n.s”.

From taxonomy to phenotype assessment, the METAGENassist tool was used. Variations in metabolic composition of the saline and non-saline sample were recorded (Figure 10). In the Non-Saline sample group, the abundance of functions such as Nitrogen fixation, Sulfur oxidation, Sulfur metabolism, and xylan degradation were predominant. On the other hand, Nitrite reduction, carbon fixation, Methane oxidation, and Ammonia oxidation were predominant functions in the saline group. From figure 11 it can also observe that, the relative abundance of anaerobic microbes are noticeably higher in saline samples (36.4%) then non-saline samples (29.3)

**Figure 10:**
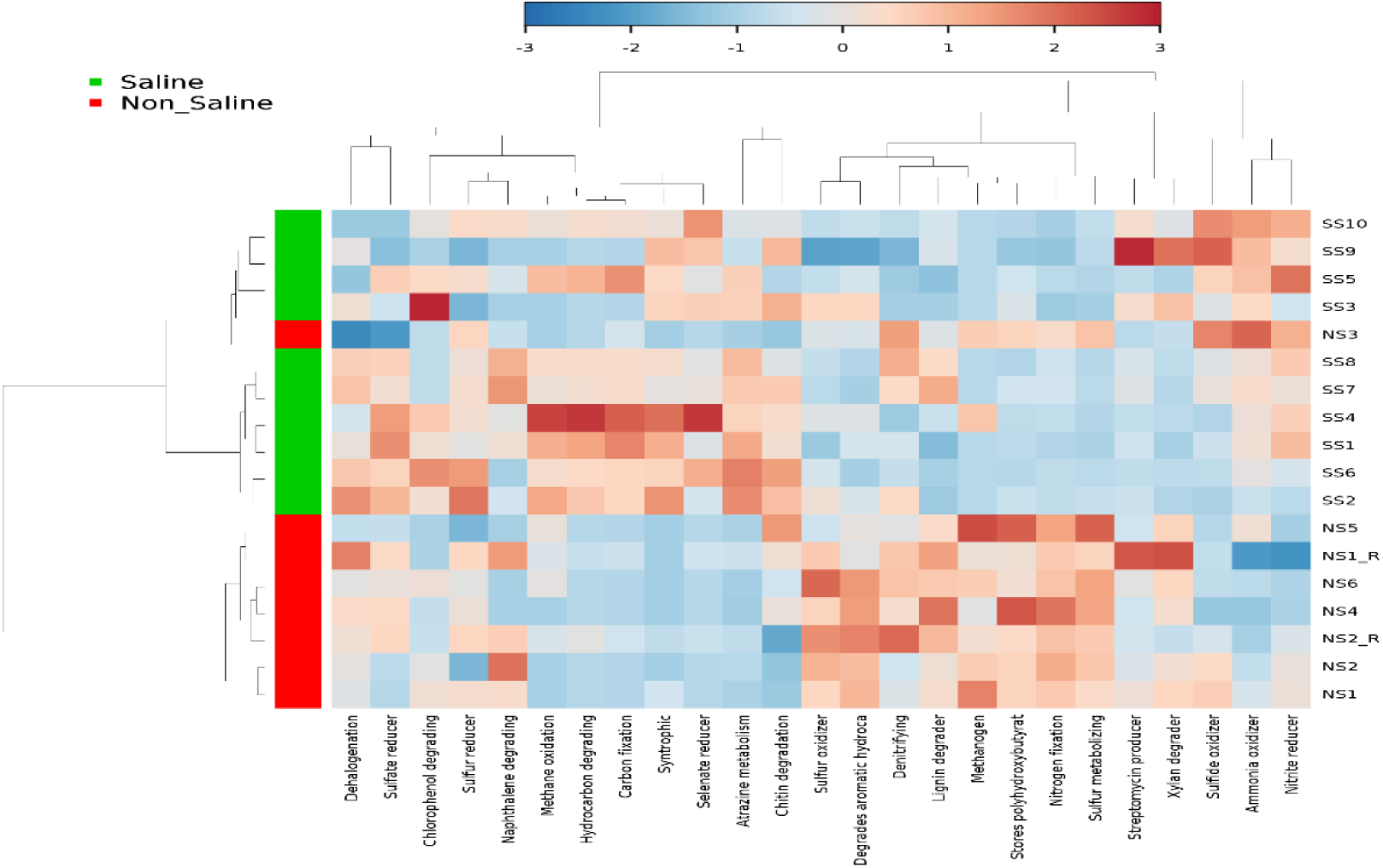
Heatmap represents functional profiles of Saline and non-saline sample groups. With Ward’s Minimum Variance Clustering Method, METAGENassist produced the heatmap With a value ranging from -2 (lowest abundance) to 4 (highest abundance). Blue cells represent less abundance, while red cells represent more abundant patterns.

**Figure 11:**
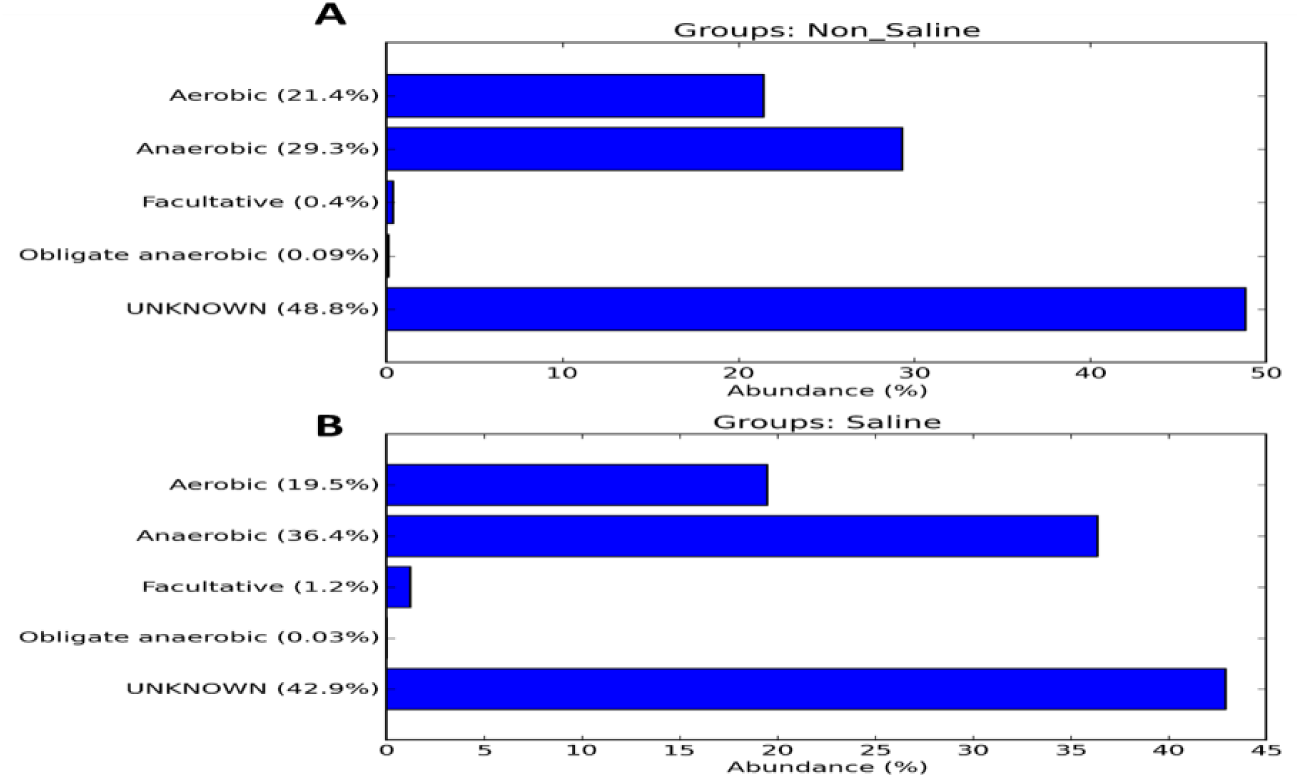
Distribution of Aerobic and anaerobic microbes in saline and non-saline soil samples represented in bar graph.

## 4. Discussion

Here in this study, the microbial community structure, and functional potential in saline and non-saline agricultural soils from coastal areas of Kuakata, Bangladesh have been analyzed and reported. Soil salinity is an important factor that decreases microbial diversity, affects the cycling of nutrients and negatively impacts the activities of plants ^13^. Fields where saline soil samples were collected had experienced total crop loss, as indicated by the presence of yellowing to blackish discoloration, which pointed to severe nutrient deficiencies and the likelihood of toxic ion accumulation ^1^. A key to addressing agricultural loss in coastal regions lies in understanding how soil salinity disrupts the abundance and functional contributions of microbial populations.

Microbial diversity was overall lower in saline soils compared to non-saline soils, although no significant differences were shown for some alpha diversity indices (Shannon, Chao1) in this study. This supports previous findings that high-salt environments pose selective pressures on microbial communities by selecting for halotolerant and especially halophilic taxa while reducing overall taxonomic richness ^14^. Beta diversity analysis indicated contrasting structures of the microbial community for saline and non-saline soils confirming the effect of salinity in the microbial composition ^9^.

Proteobacteria and Chloroflexi dominated in both saline and non-saline soils but varied in relative abundance. The relative abundance of Desulfobacterota and Deferrisomatota were significantly enriched in saline soils, presumably owing to their roles as sulfate-reducing and in metal cycling, which is critical for microbial survival and functioning at high salinity and anoxic conditions ^15^. Nonetheless, the enrichment of these taxa may be partly driven by waterlogging, or the availability of organic carbon which could play a common role in the cycling of available nutrients. In contrast, Myxococcota, Methylomirabilota and Verrucomicrobiota, which take part in organic matter degradation, nitrogen cycling and soil fertility maintenance were more abundant in non-saline soils ^16^ ^17^. These results indicate functional divergence between saline and non-saline soils.

At the genus level, functional key microbes presented opposite distributions between saline and non-saline soils. A much higher abundance of Desulfobacca, Pseudomonas and Thioalkalispira-Sulfurivermis in saline soils points to their adaptations to high-sulfate, nitrogen-limited habitats ^18^. As these genera are associated with denitrification, sulfate reduction, and sulfur metabolism, they are frequently stimulated during salt stress conditions ^19^ ^15^. In contrast, Sideroxydans, Candidatus Nitrotoga and Candidatus Solibacter were more abundant in non saline soils reflecting their involvement in iron oxidation, nitrification and organic matter mineralization essential for the health and balance of the soil ecosystem ^20^ ^21^ ^22^. Such differences emphasize functional adaptations to salinity.

This study found that anaerobic microbial abundance was higher in saline soils (36.4%), than that in non-saline soils (29.3%), which is one of the important discoveries. Anaerobic metabolism is favored under saline conditions, which are often found together with waterlogging, and this transition causes several deleterious consequences, including reduced soil fertility and impaired plant health. Saline soils often retain higher amounts of water which subsequently lowers oxygen diffusion and creates hypoxic/anoxic conditions ^23^. This favors anaerobic over aerobic microbes, disrupting key soil functions by suppressing decomposers and nitrifiers.

For example, Desulfobacterota (sulfate-reducing bacteria, SRB) were in higher abundance in saline soils. These microbes turn sulfate (SO₄²⁻) to hydrogen sulfide (H₂S), a compound that impacts plant root function and microbial respiration ^15^. Although the accumulation of H₂S is contingent upon the interplay between sulfate reduction and sulfide oxidation. Although sulfur-oxidizing bacteria are capable of detoxifying H₂S, these processes are frequently constrained in waterlogged, salty soils.

Saline soils with higher anaerobic microbial abundance are indicative of increased denitrification of nitrate (NO₃⁻) into nitrogen gases (N₂, N₂O), leading to less bioavailable nitrogen for plants ^24^. Alternative pathways such as dissimilatory nitrate reduction to ammonium (DNRA) may take place under comparable conditions and may be partly offset nitrogen loss. The extent to which these processes are important in saline soils needs to be explored.

Such decrease in aerobic decomposers (e.g., Sideroxydans and Candidatus Solibacter) in saline soils indicates a decreased capacity for providing organic matter degradation and nutrient recycling.

The increased anaerobic microbial abundance in saline soils provides strong evidence that high salinity, often coupled with waterlogging, promotes anoxic, nutrient-depleted conditions that inhibit plant growth ^25^ ^26^. The dominance of sulfate reducers and denitrifiers in saline soils suggests that these soils are prone to nitrogen loss, toxic gas accumulation (H₂S), and reduced organic matter decomposition ^27^ ^28^. In contrast, non-saline soils maintain a healthier balance between aerobic and anaerobic processes, supporting efficient nutrient cycling and plant-microbe interactions.

## Conclusion

This study highlights how salinity-induced microbial shifts, often leading to a breakdown in nutrient cycling and soil fertility, contributing to crop failure and land degradation in coastal agricultural fields. The higher anaerobic microbial abundance in saline soils emphasise the need for microbial restoration strategies to sustain soil health and agricultural productivity in salt-affected areas. However, the effectiveness of these strategies must be validated through long-term field trials, and region-specific studies are needed to account for variations in soil type, climate, and land-use history.

## Reference

1. Shrivastava P, Kumar R. Soil salinity: A serious environmental issue and plant growth promoting bacteria as one of the tools for its alleviation. Saudi J Biol Sci. 2015;22(2):123–131. doi:10.1016/j.sjbs.2014.12.001

2. Butcher K, Wick AF, DeSutter T, Chatterjee A, Harmon J. Soil Salinity: A Threat to Global Food Security. Agron J. 2016;108(6):2189–2200. doi:10.2134/agronj2016.06.0368

3. Keller LP, McCarthy GJ, Richardson JL. Mineralogy and Stability of Soil Evaporites in North Dakota. Soil Sci Soc Am J. 1986;50(4):1069–1071. doi:10.2136/sssaj1986.03615995005000040047x

4. Mukhopadhyay R, Sarkar B, Jat HS, Sharma PC, Bolan NS. Soil salinity under climate change: Challenges for sustainable agriculture and food security. J Environ Manage. 2021;280:111736. doi:10.1016/j.jenvman.2020.111736

5. Jiao S, Peng Z, Qi J, Gao J, Wei G. Linking Bacterial-Fungal Relationships to Microbial Diversity and Soil Nutrient Cycling. Stegen JC, ed. mSystems. 2021;6(2). doi:10.1128/msystems.01052-20

6. AMATO M, LADD J. Application of the ninhydrin-reactive N assay for microbial biomass in acid soils. Soil Biol Biochem. 1994;26(9):1109–1115. doi:10.1016/0038-0717(94)90132-5

7. ANDERSON JPE, DOMSCH KH. QUANTITIES OF PLANT NUTRIENTS IN THE MICROBIAL BIOMASS OF SELECTED SOILS. Soil Sci. 1980;130(4):211–216. doi:10.1097/00010694-198010000-00008

8. Selvakumar G, Shagol CC, Kim K, Han S, Sa T. Spore associated bacteria regulates maize root K+/Na+ ion homeostasis to promote salinity tolerance during arbuscular mycorrhizal symbiosis. BMC Plant Biol. 2018;18(1):109. doi:10.1186/s12870-018-1317-2

9. Lozupone CA, Knight R. Global patterns in bacterial diversity. Proc Natl Acad Sci. 2007;104(27):11436–11440. doi:10.1073/pnas.0611525104

10. Yan N, Marschner P, Cao W, Zuo C, Qin W. Influence of salinity and water content on soil microorganisms. Int Soil Water Conserv Res. 2015;3(4):316–323. doi:10.1016/j.iswcr.2015.11.003

11. Ejaz MR, Badr K, Hassan ZU, Al-Thani R, Jaoua S. Metagenomic approaches and opportunities in arid soil research. Sci Total Environ. 2024;953:176173. doi:10.1016/j.scitotenv.2024.176173

12. Dasgupta S, Hossain MM, Huq M, Wheeler D. Climate change and soil salinity: The case of coastal Bangladesh. Ambio. 2015;44(8):815–826. doi:10.1007/s13280-015-0681-5

13. Rath KM, Rousk J. Salt effects on the soil microbial decomposer community and their role in organic carbon cycling: A review. Soil Biol Biochem. 2015;81:108–123. doi:10.1016/j.soilbio.2014.11.001

14. Zhang G, Bai J, Zhai Y, et al. Microbial diversity and functions in saline soils: A review from a biogeochemical perspective. J Adv Res. 2024;59:129–140. doi:10.1016/j.jare.2023.06.015

15. Oren A. Thermodynamic limits to microbial life at high salt concentrations. Environ Microbiol. 2011;13(8):1908–1923. doi:10.1111/j.1462-2920.2010.02365.x

16. Rakitin AL, Kulichevskaya IS, Beletsky A V., Mardanov A V., Dedysh SN, Ravin N V. Verrucomicrobia of the Family Chthoniobacteraceae Participate in Xylan Degradation in Boreal Peat Soils. Microorganisms. 2024;12(11):2271. doi:10.3390/microorganisms12112271

17. Lan J, Wang S, Wang J, Qi X, Long Q, Huang M. The Shift of Soil Bacterial Community After Afforestation Influence Soil Organic Carbon and Aggregate Stability in Karst Region. Front Microbiol. 2022;13. doi:10.3389/fmicb.2022.901126

18. Foti M, Sorokin DY, Lomans B, et al. Diversity, Activity, and Abundance of Sulfate-Reducing Bacteria in Saline and Hypersaline Soda Lakes. Appl Environ Microbiol. 2007;73(7):2093–2100. doi:10.1128/AEM.02622-06

19. Wang T, Wang H, Ran X, Wang Y. Salt stimulates sulfide−driven autotrophic denitrification: Microbial network and metagenomics analyses. Water Res. 2024;257:121742. doi:10.1016/j.watres.2024.121742

20. James MT, Farrisi ST, Shah S, Shah V. Identification of Major Organisms Involved in Nutritional Ecosystem in the Acidic Soil From Pennsylvania, USA. Front Environ Sci. 2022;10. doi:10.3389/fenvs.2022.766302

21. Lantz MA, Boddicker AM, Kain MP, Berg OMC, Wham CD, Mosier AC. Physiology of the Nitrite-Oxidizing Bacterium Candidatus Nitrotoga sp. CP45 Enriched From a Colorado River. Front Microbiol. 2021;12. doi:10.3389/fmicb.2021.709371

22. Zhou N, Keffer JL, Polson SW, Chan CS. Unraveling Fe(II)-Oxidizing Mechanisms in a Facultative Fe(II) Oxidizer, Sideroxydans lithotrophicus Strain ES-1, via Culturing, Transcriptomics, and Reverse Transcription-Quantitative PCR. Buan NR, ed. Appl Environ Microbiol. 2022;88(2). doi:10.1128/AEM.01595-21

23. Barrett-Lennard EG. The interaction between waterlogging and salinity in higher plants: causes, consequences and implications. Plant Soil. 2003;253(1):35–54. doi:10.1023/A:1024574622669

24. Huang D, Li X, Luo X. Response of Nitrifier and Denitrifier Abundance to Salinity Gradients in Agricultural Soils at the Yellow River Estuary. Agronomy. 2022;12(7):1642. doi:10.3390/agronomy12071642

25. Długosz J, Piotrowska-Długosz A, Breza-Boruta B. The effect of differences in soil water content on microbial and enzymatic properties across the soil profiles. Ecohydrol Hydrobiol. 2024;24(3):547–556. doi:10.1016/j.ecohyd.2023.06.010

26. Kumar A, Singh S, Gaurav AK, Srivastava S, Verma JP. Plant Growth-Promoting Bacteria: Biological Tools for the Mitigation of Salinity Stress in Plants. Front Microbiol. 2020;11. doi:10.3389/fmicb.2020.01216

27. Akhtar M, Hussain F, Ashraf MY, Qureshi TM, Akhter J, Awan AR. Influence of Salinity on Nitrogen Transformations in Soil. Commun Soil Sci Plant Anal. 2012;43(12):1674–1683. doi:10.1080/00103624.2012.681738

28. Xin Y, Zhang H, Wu Y, et al. Salinization of coastal saline-alkali soil might enhance H2S release by affecting H2S-related bacterial communities. Appl Soil Ecol. 2023;184:104787. doi:10.1016/j.apsoil.2022.104787

